# *De novo* genome assembly and comparative genomics for the colonial ascidian *Botrylloides violaceus*

**DOI:** 10.1101/2023.07.10.548363

**Authors:** Jack T. Sumner, Cassidy L. Andrasz, Christine A. Johnson, Sarah Wax, Paul Anderson, Elena L. Keeling, Jean M. Davidson

## Abstract

Ascidians have the potential to reveal fundamental biological insights related to coloniality, regeneration, immune function, and the evolution of these traits. This study implements a hybrid assembly technique to produce a genome assembly and annotation for the botryllid ascidian, *Botrylloides violaceus*. A hybrid genome assembly was produced using Illumina, Inc. short and Oxford Nanopore Technologies long-read sequencing technologies. The resulting assembly is comprised of 831 contigs, has a total length of 121 Mbp, N50 of 1 Mbp and a BUSCO score of 96.1%. Genome annotation identified 13K protein-coding genes. Comparative genomic analysis with other tunicates reveals patterns of conservation and divergence within orthologous gene families even among closely related species. Characterization of the Wnt gene family, encoding signaling ligands involved in development and regeneration, reveals conserved patterns of subfamily presence and gene copy number among botryllids. This supports the use of genomic data from non-model organisms in the investigation of biological phenomena.

## Introduction

High-quality reference genomes of core model organisms have transformed modern biology (Bonini and Berger 2017); however, scarce genomics research on non-model species has historically limited insights from across the tree of life. Genomics technologies have become more amenable to discoveries in non-model organism biology with the rise of high throughput sequencing (da Fonseca et al. 2016; Da Fonseca et al. 2020; Etherington et al. 2020; Whitacre et al. 2017). Genome sequence data from a more diverse range of organisms will enable deeper understanding of both patterns of genome evolution and the genetic underpinnings of diverse biological phenomena.

The hybrid *de novo* genome assembly method has enabled cost-effective genome projects to improve existing reference genomes and to assemble novel drafts (Mostovoy *et al*. 2016; Xing *et al*. 2019; Wallberg et al. 2019; Tan et al. 2018; Jiang et al. 2019). This method integrates multiple technologies to produce assemblies of greater quality by complementing the strengths and weaknesses of each approach (Utturkar et al. 2014). A common hybrid assembly method combines high-coverage, short-read and low-coverage, long-read sequencing data to more accurately resolve complex repetitive regions (Tan et al. 2018; Utturkar et al. 2014; Zimin *et al*. 2013). This genome assembly technique provides low-cost yet functional sequence data. We define “functional” as sufficient to uncover novel biological insights, especially when compared to closely related species. Here we use this approach for a colonial ascidian, a member of the tunicate lineage.

Tunicates are a diverse group of marine invertebrate chordates that have been studied in the context of developmental biology, evolutionary genomics, and regenerative medicine. As the closest extant relative of vertebrates, tunicates are key to understanding the rise of vertebrate traits and the history of chordate evolution (Delsuc *et al*. 2006). Tunicate genomes are rapidly evolving and diverse (Berna and Alvarez-Valin 2014; Denoeud *et al*. 2010). Several published genomes from across the three classes of tunicates have facilitated the rise of comparative and functional genomics in this lineage (Bliznina et al. 2021; Blanchoud et al. 2018; DeBiasse et al. 2020; Dehal et al. 2002; Jue et al. 2016; Satou et al. 2019). These have revealed substantial genomic changes such as a high degree of gene loss (Dehal et al. 2002; Berna and Alvarez-Valin 2014) and unusually fast sequence evolution (Denoeud et al. 2010; Berna and Alvarez-Valin 2014). In addition, massive reorganization events have diverged their global genomic architecture from other metazoans; only faint syntenic relationships are observed even between higher taxa of tunicates (Denoeud et al. 2010). These high rates of genomic reorganization and compaction make tunicate genomes uncommonly plastic (Berna and Alvarez-Valin 2014; Denoeud *et al*. 2010), making them a valuable system for further insights about genome evolution.

Tunicate development and life histories are also highly variable; coloniality has evolved multiple times and regenerative ability varies (Lemaire 2011). Colonial ascidians, containing many discrete bodies called zooids that share a peripheral vasculature, have acquired asexual budding that mediates colony growth (Zeng *et al*. 2006; Alié et al. 2018; Berrill 1941; Watanabe and Newberry 1976). This innovation may be associated with extensive regenerative capacity; until recently, colonial ascidians in the botryllid clade were the only known chordates capable of whole-body regeneration (WBR) (Blanchoud et al. 2018; Rinkevich et al. 1995; Kassmer et al. 2020). Botryllids, consisting of the *Botrylloides* and *Botryllus* genera, are capable of WBR but show differences in its regulation (Rinkevich et al. 2007; Brown et al. 2009; Voskoboynik et al. 2007). The recent discovery of a solitary ascidian capable of regenerating all body structures from an amputated fragment (Gordon *et al*. 2021) reveals additional complexity in the evolution of regenerative abilities and further emphasizes the potential of ascidians as model organisms for studies of regeneration and stem cell biology.

Regenerative abilities across species are diverse and differ in phylogenetic distribution, with evidence for losses in various lineages (Bely 2010; Lai and Aboobaker 2018). Regeneration in vertebrates is generally more limited, including muscle (Zullo *et al*. 2020), liver (Delgado-Coello 2021), or distal structures such as limbs or tails in various fish and amphibian species (Darnet *et al*. 2019; Ferrario et al. 2020; Yokoyama 2008). Historically the best models for whole-body regeneration have been planarians (*Platyhelminthes*) and Cnidaria such as *Hydra* (Reddien and Alvarado 2004; Reddy *et al*. 2019), but there is increasing recognition of the wide range of regenerative abilities across species (Lai and Aboobaker 2018; Ferrario et al. 2020; Edgar et al. 2021; Mokalled and Poss 2018).

One fundamental question is the extent to which the genetic pathways underlying regeneration are conserved. The Wnt signaling pathway has been implicated in the regulation of stem cells and regeneration across many taxa, including ascidians (Kassmer et al. 2020; Borisenko et al. 2021; Clevers et al. 2014; Garcia et al. 2018; Leucht et al. 2019; Nusse and Clevers 2017; Zondag et al. 2016). The Wnt gene family encodes secreted glycoprotein ligands that initiate signaling pathways leading to changes in transcription associated with cell fate and proliferation, as well as cell and tissue organization (Willert and Nusse 2012; Anthony et al. 2020; Niehrs 2012). Evolution of Wnt genes, including changes in gene copy number, may play a role in varying regenerative capabilities as well as other aspects of development (Somorjai et al. 2018; Martí-Solans et al. 2021).

Although scarce genomic data has historically limited molecular investigations of botryllid biology, published genomes from the two sister genera, *Botrylloides leachii* and *Botryllus schlosseri*, are now available (Blanchoud et al. 2018; Voskoboynik et al. 2013).

Additional ascidian genome sequences will help reveal lineage-specific diversification of genes, such as Wnts, and provide the basis for further investigation of potential roles in regulating regeneration. Undergraduate students carried out the entire genome project, from obtaining local samples, optimizing DNA extraction, constructing libraries and carrying out sequencing, as well as all subsequent analysis.

This study used Illumina, Inc. short reads and Oxford Nanopore Technologies long reads to produce the first draft hybrid assembly genome sequence of the colonial ascidian *Botrylloides violaceus* (Fig. 1). Annotation of the genome allowed initial comparative genomics with other tunicates and a comprehensive survey of Wnt genes. This genome will allow further exploration of genome evolution and mechanisms involved in coloniality and regeneration.

**Figure 1.**
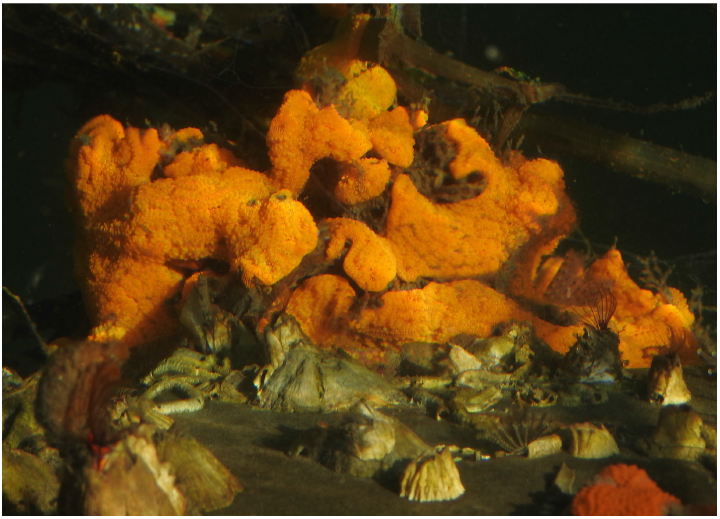
*Botrylloides violaceus* imaged in Morro Bay, CA.

## Materials and methods

### Sample Acquisition

Wild *B. violaceus* colonies were collected from Morro Bay, California (CDFW permit GM-190280002-19028) and subsequently cleared of visible detritus with forceps and brush. Colonies were cultivated and starved in filtered (0.22 µm) seawater at 10°C for approximately two days to reduce potential microbial contamination from the gut. Approximately 100 µL of blood was extracted from a colony using a 1 mL needle and syringe. DNA for Illumina sequencing was isolated using a Qiagen DNeasy Blood & Tissue Kit according to manufacturer instructions. Initial ONT sequencing using the same DNA used for Illumina sequencing produced low quality sequencing data (data not shown). From another colony, enriched blood was acquired by macerating 1 g of colony with a syringe plunger against a 40 µm sieve (Falcon™ Cell Strainer - Fisher Science) as previously described (Rosental *et al*. 2018). Over 5 µg of DNA was extracted per 200 µL enriched blood using the Zymo Quick-DNA HMW MagBead Kit. Manufacturer’s instructions were followed except for increasing the volume of magnetic beads to 50 µL total. Samples were visually assessed via gel electrophoresis and quantified with Qubit Fluorometric Quantitation (ThermoFisher, Inc.).

### Whole Genome Sequencing and Processing

#### Illumina Sequencing

DNA was processed using the Nextera XT v3 Library Preparation Kit (Illumina, San Diego, Ca, USA) to construct two libraries for short-read sequencing. 6.3 Gbp of data in 41.8 million reads (2 × 150 cycles) were produced using the MiniSeq System (Illumina, San Diego, Ca, USA). Illumina short reads were quality checked using FastQC v0.11 (http://www.bioinformatics.babraham.ac.uk/projects/fastqc). Forward and reverse reads from each library were concatenated into one pair of fastq files. Adapter sequences and poor quality data were removed from reads with Trimmomatic v0.36 (ILLUMINACLIP:2:30:10, LEADING:5, TRAILING:5, SLIDINGWINDOW:4:20, MINLEN:50) (Bolger *et al*. 2014). The Nextera XT adapter sequence (5’-CTGTCTCTTATACACATCT-3’) was supplied for trimming in fasta format. Bacterial, archaeal and viral environmental contaminants were deconvoluted from presumptive *B. violaceus* sequence data using Kraken v1.1.1 (Wood and Salzberg 2014) with the MiniKraken 8GB database. This processing culminated in approximately 37.4 million “clean” paired-end reads.

#### ONT Sequencing

Libraries for ONT sequencing were constructed using SQK-RAD004 Rapid Sequencing Kit (Oxford Nanopore Technologies, Oxford, UK) following the manufacturer’s instructions with the exception that the DNA mass input was increased to approximately 1 µg. ONT libraries were sequenced using the MinIon system (Oxford Nanopore, Oxford, UK) to produce 4.08 Gbp of sequencing data with an N50 of 9,585 bp. Nanopore long reads were basecalled using Guppy v4.0.15 (– config dna_r9.4.1_450bps_fast.cfg) (Oxford Nanopore Technologies). Reads were then quality filtered to include those with quality scores greater than or equal to seven (approximately 85% basecall accuracy) using NanoFilt v2.7.1 (De Coster et al. 2018). Filtering resulted in 3.56 Gbp of long-read data, with a read length N50 of 9,799 bp.

### Genome Assembly and Annotation

#### Genome Assembly

Standard short-read, long-read, and hybrid assembly approaches were performed on appropriate data. We employed the MaSuRCA v3.4.2 (Zimin et al. 2017) algorithm for hybrid *de novo* assembly of both reads types with the Flye assembler for final assembly of mega reads, rather than previous CABOG versions, for improved efficacy and efficiency. Short and long-read only assemblies were produced with ABySS v2.0.2 (Jackman *et al*. 2017) and Flye v2.8.1 (Kolmogorov et al. 2019), respectively. To estimate completeness, BUSCO analysis (Fig. 2) was performed on each method and key genome assembly statistics were calculated with QUAST (see Genome Statistics methods).

**Figure 2.**
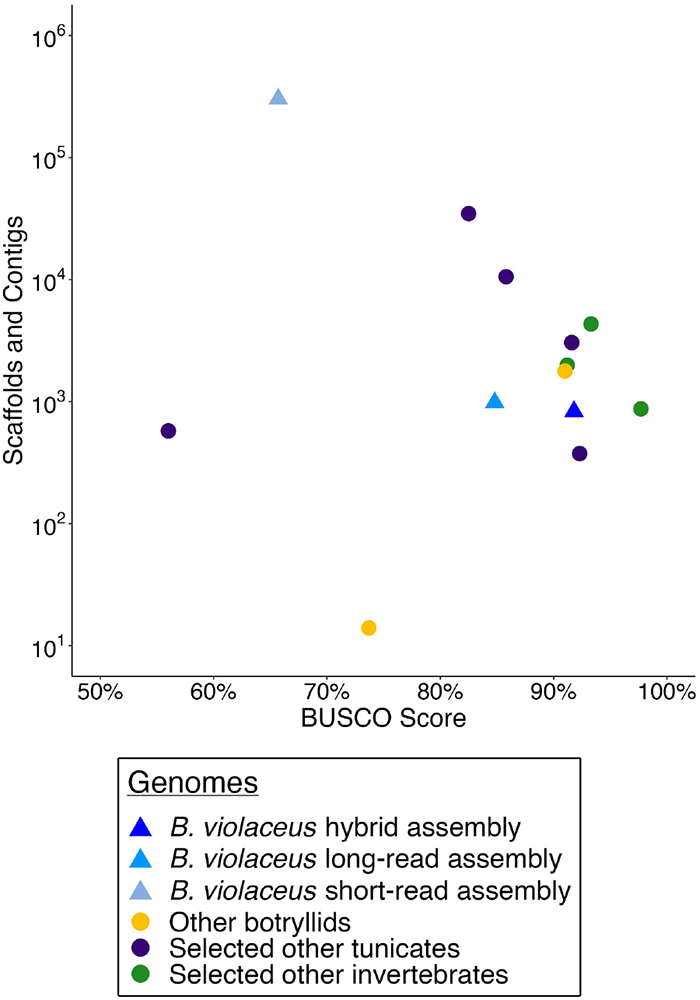
Using BUSCO score and the total number of scaffolds and contigs in genome assemblies to assess their quality and completeness. *B. violaceus* assemblies are compared to genomes of other botryllids (*B. leachii, B. schlosseri*), other selected tuni-cates (*O. dioica, P. fumigata, C. savignyi, H. roretzi, M. oculata*), and other selected invertebrates (*S. purpuratus, A. californica, H. robusta*). Triangular points are *B. violaceus* assemblies color coded by assembly type, while circular points represent other genomes color coded by grouping. BUSCO scores are the percentage of complete BUSCOs based on the Metazoa dataset.

#### Genome Annotation

Annotation of the *B. violaceus* hybrid assembly was completed using the Genome Sequence Annotation Server (GenSAS) v6.0 (Humann et al. 2019). GenSAS allowed all listed annotation analyses to be streamlined into one pipeline. Prior to annotation, the hybrid assembly was filtered to only contain scaffolds and contigs 1,000 bp or greater, which resulted in removing 101 contigs (filtered assembly length 120,854,445 bp). Removing these contigs did not change the BUSCO score. Since this is a *de novo* annotation, no prior *B. violaceus* data or results were uploaded to assist in annotation. However, since the sister species *B. leachii* has an annotated genome assembly and transcriptome, the *B. leachii* transcripts and were obtained from ANISEED (Brozovic et al. 2018; Tassy et al. 2010) and uploaded to GenSAS to use as evidence in downstream programs. RepeatMasker v4.0.7 (http://www.repeatmasker.org) was used with default parameters, NCBI RMBlast v2.2.27 search engine and another well-studied tunicate, *Ciona intestinalis*, as the pre-masked DNA source to aid in finding repetitive regions in the *B. violaceus* genome. RepeatModeler v1.0.11 (http://www.repeatmasker.org/RepeatModeler/) was used to find repetitive elements *de novo* in the hybrid assembly. The results from RepeatMasker and RepeatModeler were combined into a consensus masked genome that was used in downstream annotation (GenSAS built-in masked consensus tool). For structural annotation of the genome, *B. leachii* transcripts were aligned to the *B. violaceus* hybrid assembly using blastn v2.7.1 (Camacho *et al*. 2009) with an e-value cutoff of 1e-30 and all other parameters left default. *Ab initio* gene predictions were made using GeneMark-ES v4.38 (Lomsadze et al. 2005) with the max_contig set to 6,000,000 bp, larger than the hybrid assembly’s longest contig as to not split the contigs, and the min_contig set to 1,000 bp to include all contigs in the training of the algorithm; all other parameters were left default. EVidenceModeler v1.1.1 (Haas et al. 2008) combined the *B. leachii* transcript alignments from blastn and the *ab initio* gene predictions from GeneMark-ES into one consensus structural annotation using the default weights of 10 for the transcripts and 1 for the *ab initio* gene predictions. This consensus of gene predictions was assigned as the Official Gene Set (OGS) of which functional annotation would take place using InterProScan v5.44-79.0 (Jones *et al*. 2014). Additional functional annotation of structurally annotated proteins was conducted outside of GenSAS to create an EggNOG-based annotation using eggNOG-mapper v1.0.3 (Huerta-Cepas *et al*. 2017).

#### Genome Statistics

Genome statistics including number of contigs or scaffolds, N50, and total length were obtained using the Quality Assessment Tool for genome assemblies, QUAST v5.0.2 (Gurevich *et al*. 2013). Benchmarking Universal Single-Copy Orthologs (BUSCO) v5.1.3 assessment (Manni *et al*. 2021) was used to estimate assembly completion using the eukaryota_odb10 (Eukaryota) and metazoa_odb10 (Metazoa) datasets compiled from OrthoDB v10 (Kriventseva *et al*. 2019). BUSCO assessments were completed independently for each genome using the genome mode. Furthermore, analysis of the annotation of the *B. violaceus* hybrid assembly was completed using BUSCO in protein mode using all protein sequences from the annotation.

### Comparative Genomics

#### Acquisition of Relevant Genomic Reference Data

Several genomes were acquired for comparative genomics analysis. Tunicate genomes including *B. leachii, B. schlosseri, H. aurantium, H. roretzi, M. occidentalis, M. oculata, P. mammilata, P. fumigata, C. savignyi*, and *O. dioica* were acquired from ANISEED (Brozovic *et al*. 2018; Tassy *et al*. 2010). Selected other invertebrate genomes used to assess the quality of the *B. violaceus* hybrid assembly included *S. purpuratus, californica*, and *H. robusta* and have RefSeq accession numbers of GCF_000002235.5, GCF_000002075.1, and GCF_000326865.1, respectively.

#### OrthoFinder Analysis

To better understand the evolution of botryllid-specific traits and to distill relevant information from high-dimensional genomic data, we implemented a gene-centric comparative genomics approach. To that end, pairwise comparisons of protein-coding genes were performed to identify groups of evolutionarily conserved genes across tunicates using OrthoFinder (Emms and Kelly 2019). This high-dimensional dataset was then refined using a stepwise analytical approach that uses publicly available bioinformatics software and in-house scripts to identify copy number variation patterns and putative functions that are unique to botryllids.

The OrthoFinder algorithm was implemented to identify groups of homologous genes (orthogroups) from across the tunicate lineage. Briefly, annotated proteomes from select tunicates (i.e., *B. leachii, B. schlosseri, H. aurantium, H. roretzi, M. occidentalis, M. oculata, P. mammilata, P. fumigata, C. savignyi*, and *O. dioica*) were downloaded from ANISEED and processed, along with the putative *B. violaceus* proteome from the GenSAS annotation, using OrthoFinder v2.5.2 with default parameters. After initial orthgroups were identified, basic quality control was performed to broadly assess the reliability of these data, as described in the OrthoFinder documentation (Emms and Kelly 2019). The inferred phylogenetic tree was visualized using iTOL (Letunic and Bork 2021) to confirm its consistency with previous reports as reliable phylogenetic reconstruction is essential to downstream analyses.

To dissect botryllid-specific patterns of copy number variations, orthogroups were evaluated using a phylogenetically informed approach. Briefly, gene trees from each node of the inferred phylogeny are clustered using Orthofinder to create lineage-specific hierarchical orthogroups (HOGs) (Altenhoff *et al*. 2013). We then selected HOGs corresponding to the botryllid lineage for our remaining analysis and hereto referred to as HOGBs. Gene counts from HOGBs were used to create a species by HOGB matrix that was organized using unsupervised hierarchical clustering; this cladogram was then cut into eighteen clusters, each containing unique sets of HOGBs. The number of clusters was determined by cutting the cladogram into one to twenty clusters and computing the average silhouette score, a metric used to assess cluster quality of fit, at each point; the local maxima was found at eighteen clusters and was thus used for the remaining analysis (Supplementary Fig. S1).

#### Gene Ontology Analysis

KEGG (Kanehisa and Goto 2000) orthology terms were assigned to each *B. violaceus* gene based on our EggNOG-based annotation. Cluster enrichment was performed using the compareCluster and enrichKEGG functions in the R package clusterProfiler v4.2.1 (Wu et al. 2021). Additionally, we assessed the presence of relevant developmentally-related pathways in each cluster. HOG cluster output provided information on which *B. violaceus* gene was within each cluster, and the corresponding KEGG annotation information for that gene. Genes with KEGG terms corresponding to investigated pathways were recorded within each cluster. The following KEGG pathways were investigated: Wnt (ko04310), TGF-beta (ko04350), Hedgehog (ko04340), Notch (ko04330), ErbB/EGF (ko04012), TNF (ko04668), cytokine-cytokine receptor interaction (ko04060), and retinol metabolism (ko00830).

### Wnt Family Analysis

Annotation data were used to identify putative Wnt pathway gene orthologs in the *B. violaceus* genome. We then focused on characterizing the repertoire of Wnt subfamilies, using BLAST and multiple sequence alignment analyses.

Using the functional annotation data from InterProScan, genes with the IPR term IPR005817 (Wnt description) were aligned against the NCBI non-redundant nucleotide collection (blastn with nr/nt) and nonredundant protein sequences (blastp with nr) databases to confirm Wnt gene orthologs. Indeterminate results showing multiple putative subfamilies for a single Wnt sequence were clarified using the ANISEED built-in BLAST tool with tblastn (Brozovic et al. 2018); similarity searches were performed against two Gene Model databases, *B. leachii* (SBv3) and *B. schlosseri* (2013), with default parameters. Additional methods were required to support identification of Wnt genes 1, 11, and A in *B. violaceus* utilizing Integrative Genomics Viewer (IGV) v2.9.2 (Thorvaldsdóttir *et al*. 2013) and additional BLAST and MSA analyses. Upon manual curation and phylogenetic analysis (described below) of Wnt sequences, we identified several *B. leachii* Wnts that had greater identity to alternative subfamilies than previously described (Blanchoud *et al*. 2018) and reclassified them accordingly for our analysis (supplemental data on GitHub).

For phylogenetic analysis of Wnt subfamilies, orthologous Wnt protein sequences from twelve different species (tunicates and human) were obtained from Martí-Solans *et al*. (2021). Wnt protein sequences from *B. leachii*, obtained from ANISEED, and *B. violaceus*, identified in this study, were added to the those twelve species for comprehensive phylogenetic analysis. A custom, automated phylogeny workflow was built in NGPhylogeny (Lemoine *et al*. 2019) and was used to construct the Wnt phylogeny. The Wnt protein sequences were aligned in Clustal Omega (Sievers *et al*. 2011). The maximum likelihood phylogeny constructor PhyML v3.3 (Guindon *et al*. 2010) was used with the default parameters for amino acid data and SH-like aLRT (Shimodaira–Hasegawa-like approximate likelihood ratio test) branch supports. The phylogeny was annotated and edited in iTOL (Letunic and Bork 2021). Determination of Wnt subfamily presence and copy number in other species was based on published literature (Somorjai *et al*. 2018; Martí-Solans *et al*. 2021; Nayak *et al*. 2016). Species relationships used the current accepted phylogeny of tunicates (DeBiasse *et al*. 2020).

## Results and discussion

### Genome Assembly and Annotation

The ABySS (short-read), Flye (long-read), and MaSuRCA (hybrid) genome assemblers produced assemblies of lengths 160.1 Mbp, 122.2 Mbp, and 120.9 Mbp respectively (Table 1). Although the hybrid assembly technique did produce the smallest draft genome, it produced the most contiguous genome. The MaSuRCA hybrid assembly contains the smallest number of contigs, the greatest number of contigs over 50 Kbp, and the greatest length of largest contig. The Flye assembly had a slightly larger N50 than the MaSuRCA assembly, however, both substantially surpassed the N50 of the ABySS assembly by 1,000 fold (Table 1).Though the hybrid assembly did not produce the largest N50, it did have the largest L50 indicating a higher percentage of the hybrid assembly consists of long contigs varying in length between its N50 and largest contig (Table 1). The ABySS assembly had few contigs over the length of 1,000 bp, which indicates little realistic viability of the genome for downstream analysis. While the Flye and MaSuRCA assemblies are similar in structural statistics, the MaSuRCA hybrid assembly represents the most complete and functional draft genome for *Botrylloides violaceus* and is used as the reference genome for the remainder of the analysis on this species.

**Table 1.** Genome statistics for *B. violaceus* assemblies.

|  | **ABYSS**<br>*Short-read assembly* | **Flye**<br>*Long-read assembly* | **MaSuRCA**<br>*Hybrid assembly* |
| --- | --- | --- | --- |
| Number of contigs | 302,459 | 981 | 831 |
| Number of contigs ≥ 1,000 bp | 31,964 | 840 | 730 |
| Percent contigs ≥ 1,000 bp | 10.6 | 85.6 | 87.8 |
| Number of contigs ≥ 50,000 bp | 0 | 175 | 245 |
| Largest contig (bp) | 44,514 | 5,304,203 | 5,758,483 |
| Total length (bp) | 160,170,160 | 122,263,676 | 120,925,257 |
| N50 (bp) | 1,073 | 1,647,705 | 1,028,308 |
| L50 | 29,444 | 18 | 28 |
| Number N's per 100 kbp | 3.73 | 1.31 | 1.41 |

MaSuRCA estimated the *B. violaceus* genome size to be 139 Mb, suggesting that around 86% of the genome has been assembled (Table 2). The hybrid assembly and estimated genome size are the smallest of the botryllids where current draft genome assemblies are sized at 159Mbp for *B. leachii* and 580Mbp for *B. schlosseri*, and estimated genome sizes are 194Mbp for *B. leachii* and 725 Mbp for *schlosseri* (Blanchoud et al. 2018; Voskoboynik et al. 2013).

**Table 2.**
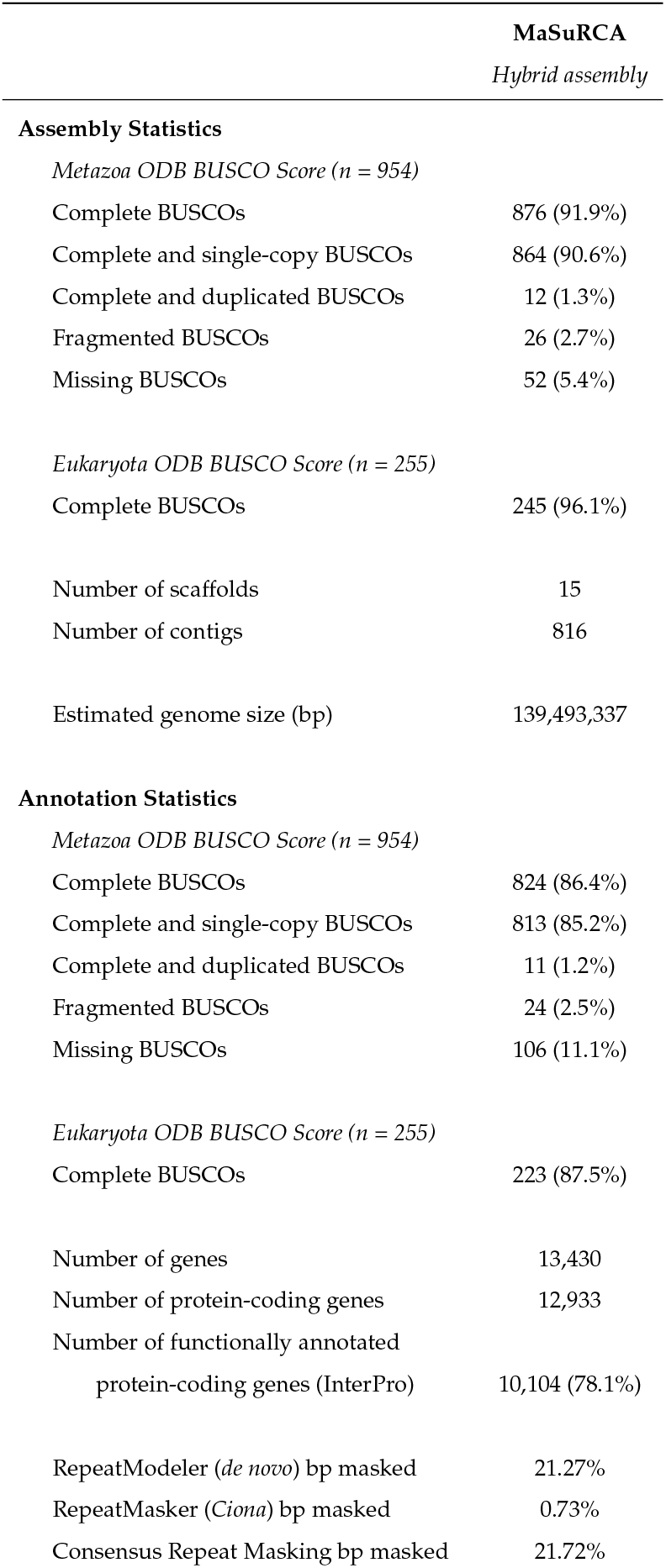
Additional hybrid assembly statistics, including relevant annotation statistics.

|  | MaSuRCA<br>Hybrid assembly |
| --- | --- |
| **Assembly Statistics** |  |
| *Metazoa ODB BUSCO Score (n = 954)* |  |
| Complete BUSCOs | 876 (91.9%) |
| Complete and single-copy BUSCOs | 864 (90.6%) |
| Complete and duplicated BUSCOs | 12 (1.3%) |
| Fragmented BUSCOs | 26 (2.7%) |
| Missing BUSCOs | 52 (5.4%) |
| *Eukaryota ODB BUSCO Score (n = 255)* |  |
| Complete BUSCOs | 245 (96.1%) |
| Number of scaffolds | 15 |
| Number of contigs | 816 |
| Estimated genome size (bp) | 139,493,337 |
| **Annotation Statistics** |  |
| *Metazoa ODB BUSCO Score (n = 954)* |  |
| Complete BUSCOs | 824 (86.4%) |
| Complete and single-copy BUSCOs | 813 (85.2%) |
| Complete and duplicated BUSCOs | 11 (1.2%) |
| Fragmented BUSCOs | 24 (2.5%) |
| Missing BUSCOs | 106 (11.1%) |
| *Eukaryota ODB BUSCO Score (n = 255)* |  |
| Complete BUSCOs | 223 (87.5%) |
| Number of genes | 13,430 |
| Number of protein-coding genes | 12,933 |
| Number of functionally annotated<br>protein-coding genes (InterPro) | 10,104 (78.1%) |
| RepeatModeler (*de novo*) bp masked | 21.27% |
| RepeatMasker (*Ciona*) bp masked | 0.73% |
| Consensus Repeat Masking bp masked | 21.72% |

BUSCO scores for the hybrid assembly show over 90% of highly conserved orthologs present from both the Metazoa and Eukaryota datasets (Table 2). To further assess the quality of the *B. violaceus* assemblies, comparisons were drawn using the other botryllid genomes (*B. leachii* and *B. schlosseri*), other tunicates, and other invertebrates (Fig. 2). BUSCO score (complete single-copy and duplicated BUSCOs) from the Metazoa dataset is used to assess genome completion and the total number of scaffolds and contigs in each genome is used to assess the contiguity and quality. Compared to all other assemblies, the *B. violaceus* hybrid assembly is in the top 25% of BUSCO scores while having an intermediate scaffold and contig number. Both the *B. violaceus* short and long-read assemblies have a lower BUSCO completeness score and greater numbers of scaffolds and contigs compared to the hybrid assembly. The other selected invertebrates tend to have higher BUSCO scores compared to tunicates, which is consistent with previous accounts of genomic loss events in the tunicate ancestor (Seo et al. 2001). Our *B. violaceus* hybrid assembly is therefore of high quality and completeness when compared to tunicates, and is comparable to other published invertebrate genomes (Fig. 2).

Annotation of the *B. violaceus* hybrid genome assembly provided sufficient predictions to use in future analyses. The GenSAS annotation of the hybrid genome assembly predicted 13.4K genes, 12.9K protein-coding genes, with 78% of protein-coding genes being functionally annotated with InterProScan. In total, 21% of the genome was masked as repetitive regions. Additionally, the protein annotation had a BUSCO score of 86% using the Metazoan database (Table 2).

Initially, annotation used evidence including transcripts from related species and gene models produced from *ab initio* gene predictors such as SNAP (7/28/2006 version) (Korf 2004) and GeneMark-ES. However, when using SNAP to predict genes based on the *Ciona intestinalis* HMM, the resulting annotation contained significant fragmentation of the genes in the OGS in comparison to orthologs in the other botryllids, *B. schlosseri* and *B. leachii*. Removing SNAP from the OGS and using only the *B. leachii* transcript alignments and GeneMark-ES predictions rectified the fragmentation of genes and significantly decreased the number of predicted genes from 23K to 13K, which is comparable to the 15K predicted genes in *B. leachii* (Blanchoud et al. 2018). Additionally, when identifying members of the Wnt family based on the final annotation, no Wnt 1 was found in the annotation despite its presence in closely related species. However, Wnt 1 was successfully located due to the 6 *De novo* genome assembly and comparative genomics for the colonial ascidian *Botrylloides violaceus* GeneMark-ES predictions. Inconsistencies in this gene annotation may be due to heavier weighting of the *B. leachii* transcripts over the *ab initio* GeneMark-ES predictions when combining evidence to support the final gene models.

### Comparative Genomics

Genomic datasets are reservoirs of rich, high-dimensional data that can be used to infer evolutionary relationships and molecular functions through the lens of comparative genomics. Yet, distilling persuasive evidence for these biological concepts remains challenging. To that end, we implemented a step-wise analytical approach to identify copy number variation patterns that are unique to botryllids (Fig. 3A-D). Pairwise comparison of protein-coding genes has identified patterns of genome evolution unique to the botryllid lineage (see methods for details). Interpreted within an evolutionary framework, we hypothesize that these botryllid-specific patterns are representative of botryllid-specific traits such as high regenerative capacity and coloniality, which thus provide a resource for experimentally disentangling these complex phenomena.

**Figure 3.**
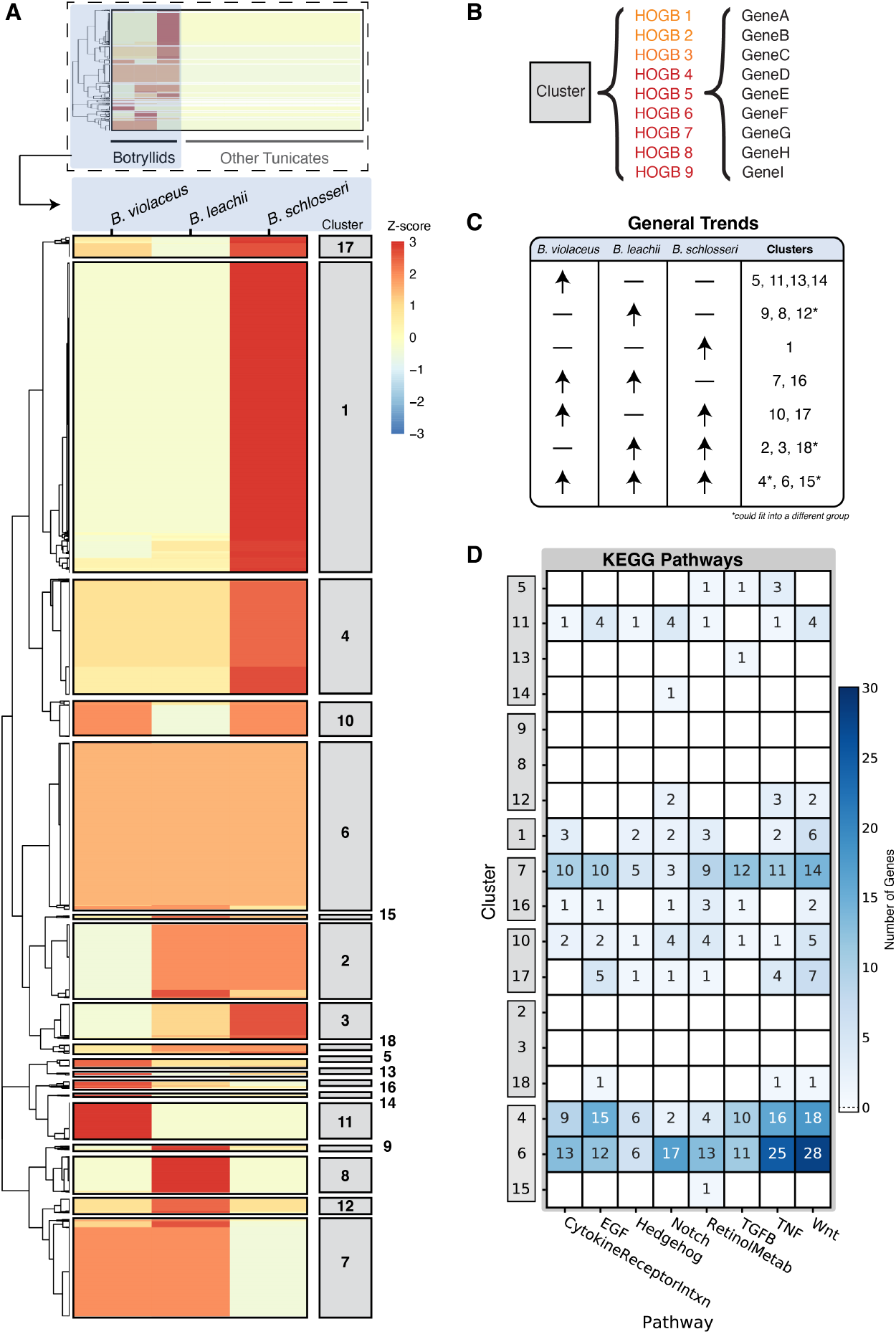
Comparative genomic analysis using hierarchical orthogroups reveals patterns of conservation and divergence in the botryllid lineage. (**A-B**) HOGBs clustered using variation in gene copy number per species. A species by HOGB copy number matrix was clustered to identify groups with similar patterns of gene family evolution in botryllids. The appropriate number of clusters (18) was chosen using the silhouette method to account for variation between clusters. (**C**) HOGB clusters binned by duplication pattern similarity. (**D**) Number of *B. violaceus* genes in each cluster that are members of developmentally relevant pathways according to KEGG annotation. Clusters ordered by general trends depicted in **C**.

Gene family evolution plays an essential role in trait acquisition (Capra et al. 2010). We thus implemented the OrthoFinder algorithm to identify groups of homologous genes (orthogroups) from across the tunicate lineage (Fig. 3A). Although some orthogroups do include genes from all tested species, not all species will be represented in all orthogroups (Emms and Kelly 2019). From the eleven tunicate proteomes used in this analysis, 229,318 genes were assigned into 21,690 orthogroups which includes 88.6% of the original gene set. The relative size of each orthogroup varies considerably and the number of homologous genes in an orthogroup ranges from 2 to 447 (mean=6.0, median=9.4). Interestingly, 10.6% of all orthogroups represent single species; this suggests that novel or retained gene lineages have expanded in single species, supporting previous reports of rapid diversification in the tunicate lineage (Berna and Alvarez-Valin 2014; Denoeud *et al*. 2010; Jue *et al*. 2016).

The degree to which species overlap in orthogroups is useful for identifying genome scale duplication and contraction events (Emms and Kelly 2019). Pairwise correlation analysis of gene counts per orthogroup reveals the extent to which species show similar duplication and loss patterns (Supplementary Fig. S2). Indeed, we observe that *O. dioica* has substantially reduced orthogroup overlap and gene counts with other tunicate lineages (Supplementary Fig. S2, Supplementary Fig. S3) which is consistent with its dramatically compacted genome (Seo *et al*. 2001). In contrast, *B. schlosseri* retains greater orthogroup overlap and gene count correlations with other botryllids than with non-botryllid tunicates (which are strikingly weak). These data support the hypothesis that secondary gene duplication events in *Botryllus* gave rise to an expanded genome after its divergence with the shared botryllid ancestor (Supplementary Fig. S2, Supplementary Fig. S3); however, they do not refute the alternative, though not mutually exclusive, hypothesis that the *B. schlosseri* genome expanded due to horizontal gene transfer (Voskoboynik *et al*. 2013).

Hierarchical orthogroups are comprised of the orthologs and in-paralogs of a specific clade. Lineage-specific hierarchical orthogroups were identified for botryllids (HOGBs) (see methods for details; Fig. 3A). To dissect botryllid-specific patterns of copy number variations, we performed cluster analysis using the gene copy number in a HOGB for each species (Fig. 3A-B). The 18 clusters of HOGBs represent changes in copy number variation among botryllids (Fig. 3A); each cluster contains unique sets of HOGBs, and each cluster represents several HOGBs which in turn include varying numbers of homologous genes (Fig. 3B). Patterns of copy number variation can be classified into seven broad categories that communicate relative changes of variation both within botryllids and between botryllids and non-botryllid tunicates (Fig. 3C). Generally, we observe that all botryllid genomes encode similar or greater HOGB gene copies than non-botryllid tunicate genomes, as expected (Fig. 3A). However, the degree to which HOGB gene copies encoded in botryllid genomes differ from non-botryllid tunicates varies in a species-dependent manner (Fig. 3C). For instance, the HOGB copy number of cluster 6 is mutually expanded in all botryllids relative to non-botryllid tunicates (Fig. 3A). This pattern suggests that gene duplications in the botryllid ancestor that were then conserved in its descendants. Similarly, cluster 16 contains HOGBs that have expanded in *Botrylloides spp*. but remained largely unchanged in *B. schlosseri*, likely indicating expansion events in the *Botrylloides* ancestor (Fig. 3A). As *Botrylloides spp*. regenerate more readily than *B. schlosseri* (Brown *et al*. 2009; Nourizadeh *et al*. 2021), these patterns may highlight genetic components involved in regulation of whole body regeneration. Overall, these data suggest that the botryllid lineage has undergone large-scale gene expansion events at multiple points in their evolutionary lineage.

In addition to overall patterns of gene losses and gains, functional annotations associated with binned clusters suggest how patterns may relate to phenotypes. For example, retinoic acid signaling has a well characterized role in regeneration and development (Rinkevich et al. 2007). Genes involved in synthesis and breakdown of retinoic acid (retinol metabolism) have been differentially lost or retained in the tunicate lineage (Blanchoud *et al*. 2018). KEGG enrichment analysis reveals that genes involved in retinol metabolism are enriched in cluster 16, representing HOGBs expanded in *Botrylloides spp*. (Supplementary Fig. S4). In addition, four paralogs of CYP26A, a P450 related gene involved in intracellular breakdown of retinoic acid, were found in the *B. violaceus* assembly and previously reported in *B. leachii* (Blanchoud *et al*. 2018). Interesting, one CYP26A ortholog found in cluster 7 was not represented in *B. schlosseri* but was present in both *Botrylloides* species, suggesting either a gene loss or gain event with potential implication for regeneration.

Other developmental signaling pathways are also important for coordination and regulation of regeneration. For example, HDAC2/3, which are effector proteins involved in Notch signaling regulation (KEGG: hsa04330), were identified in our KEGG search (Fig. 3D). HDAC2 is represented in all three botryllids (cluster 6) while *B. violaceus* HDAC3 is represented in a *Botrylloides*-enriched group (cluster 7). Inhibition of HDAC suppresses WBR in *B. leachii* (Zondag et al. 2019) and this mechanism is likely conserved in *B. violaceus*, although experimental testing is necessary to confirm. The copy number of *B. violaceus* genes putatively involved in Wnt and TNFs signaling pathways was notably high in clusters generally associated with clusters expanded in botryllids relative to non-botryllids (clusters 4, 6; Fig. 3A,D). TNFs play a major role in innate immunity, inflammation and regulation of cell death (Webster and Vucic 2020). Ascidians are useful invertebrate models of allorecognition and the evolution of the immune system in chordates (Voskoboynik et al. 2013; Franchi and Ballarin 2017). For example, over 10 orthologs of the BIRC2 gene, which negatively interacts with TNF signaling cascades, were found in the *B. violaceus* genome. These genes putatively function as inhibitors of apoptosis and several inhibitor of apoptosis genes have been implicated in botryllid regeneration and torpor (Rosner et al. 2019).

As HOGB clusters are unique to the botryllid lineage, this analysis is expected to be more sensitive to botryllid-lineage gene duplication events than it is to gene-loss events; gene-loss events that occurred early after ancestral botryllid diversification may require greater genomic resolution to detect (i.e., sampling more botryllid genomes). Furthermore, gene loss inferences may be confounded by possibly incomplete genome coverage. However, gene-loss events can be inferred when expansions are conserved between sister genera, but not within the same genus. Cluster 10 is comprised of HOGB clusters with expanded repertoires in both *B. violaceus* and *B. schlosseri*, but not in *B. leachii* (Fig. 3C). Therefore, one may infer that primary duplication events in the botryllid ancestor followed by secondary gene loss events in the *B. leachii* ancestor may have occurred. The biological significance remains elusive, although KEGG enrichment analysis reveals that genes associated with neuronal development or function may facilitate some unknown trait that may have been lost in *B. leachii* (Supplementary Fig. S4). Thus, we provide evidence that comparative genomics analysis of closely related species allows low-resolution inference of their shared ancestor’s genomic content and that interpretation within an evolutionary context is useful in generating hypotheses for future experimental testing.

### Wnt Family Analysis

Gene predictions from the hybrid assembly annotation and additional analysis allowed us to characterize the repertoire of Wnt genes found in *B. violaceus*. Characterizing Wnt genes in *B. violaceus* demonstrates the functionality of our genome and provides genetic targets for investigating the unique regenerative abilities of this organism. Comprehensive analysis of Wnt genes in the *B. violaceus* genome indicates the presence of 10 Wnt subfamilies (Fig. 4). Our final Wnt gene repertoire is corroborated by similar patterns in other tunicate genomes (Fig. 5A) (Somorjai *et al*. 2018; Martí-Solans *et al*. 2021). The initial BLAST annotations identified most, but not all, of these Wnt genes, emphasizing the necessity of additional manual annotation methods.

**Figure 4.**
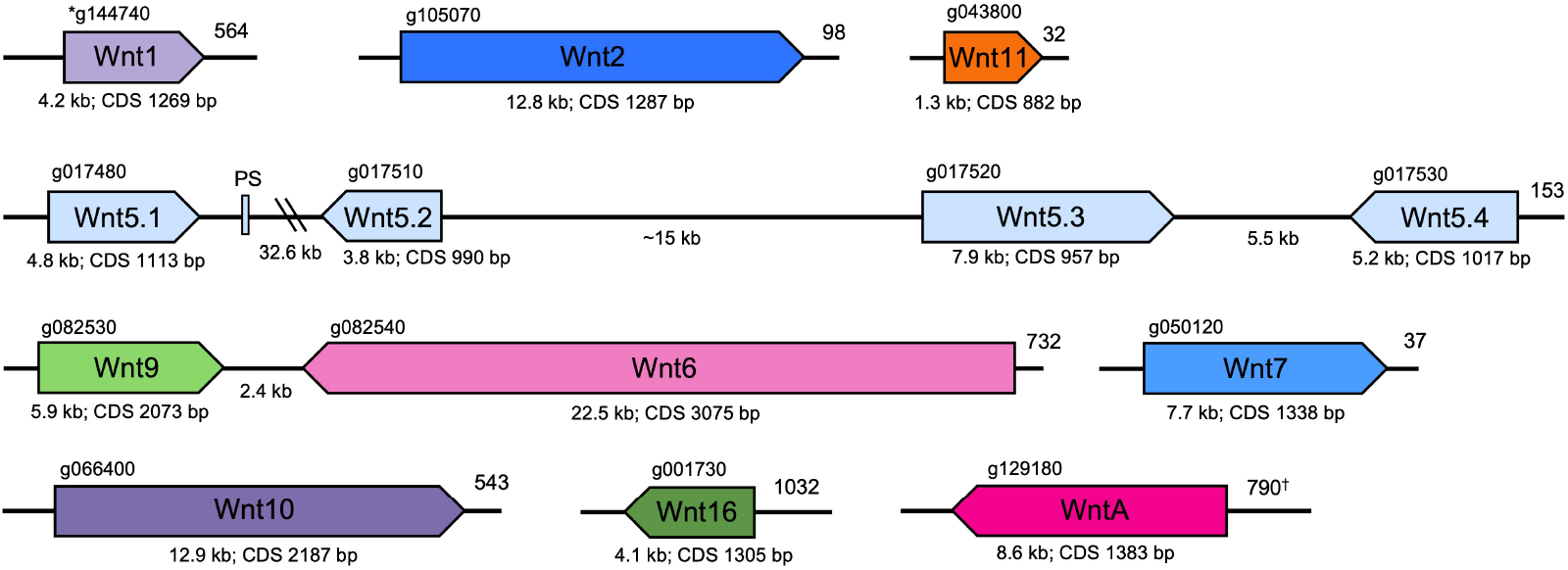
Relative position and orientation of the Wnt genes in *B. violaceus*. Distances between genes and gene sizes are to scale, with the exception of Wnt11 which was slightly enlarged for clarity. PS indicates a Wnt5 pseudogene. Gene identification number from GenSAS annotation is on top. Numbers at rightmost position of fragments refer to specific contig or scaffold^†^ in the hybrid assembly. Kb refers to genomic size; CDS bp refers to coding DNA sequence length; double-parallel lines indicate the intergenic region is > 30 kb. ^*^Wnt1 gene identification number is from GeneMark-ES gene model predictions.

**Figure 5.**
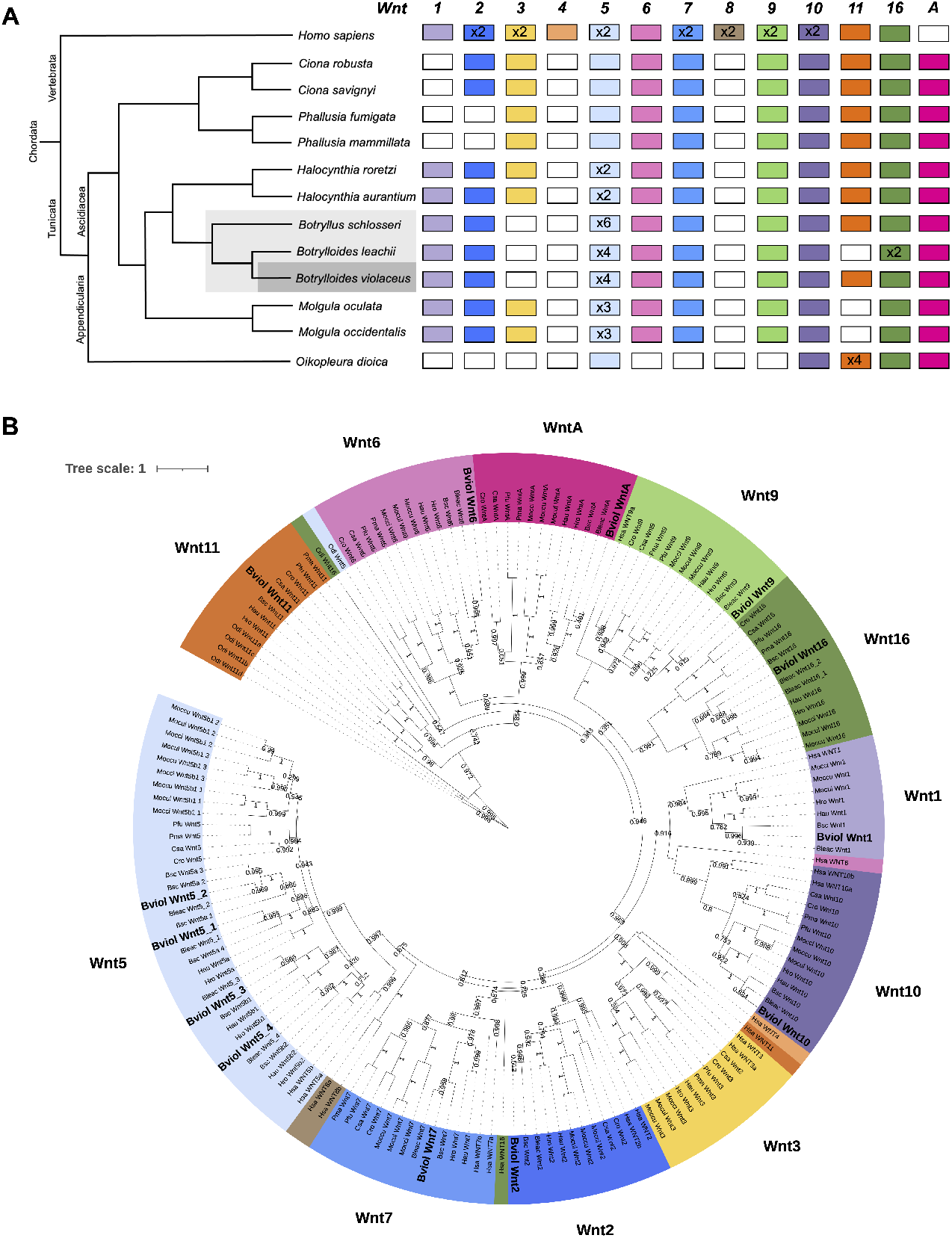
Wnt gene family analysis. (**A**) Summary of Wnt repertoire for comparative analysis of selected tunicates and humans. Coloring of the boxes indicates the presence of the indicated Wnt subfamily, while a white box indicates the absence. Total copy number of the gene is provided in the respective box. Botryllids are highlighted in grey on the phylogeny. (**B**) Maximum likelihood phylogeny of tunicate Wnt protein sequences. Branch support values derived from SH-like aLRT method. Species represented: *Botrylloides violaceus* (Bviol), *Botrylloides leachii* (Bleac), *Botryllus schlosseri* (Bsc), *Ciona savignyi* (Csa), *Ciona robusta* (Cro), *Halocynthia roretzi* (Hro), *Halocynthia aurantium* (Hau), *Mogula occulta* (Moccu), *Mogula oculata* (Mocul), *Mogula occidentalis* (Mocci), *Phallusia fumigata* (Pfu), *Phallusia mam-millata* (Pma), *Oikopleura dioica* (Odi), and *Homo sapiens* (Hsa). *O. dioica* Wnt10 was removed from the Wnt10 cluster due to long branch length in order to increase visibility of other leaves. Scale bar represents amino acid substitutions.

Our workflow for manually confirming the Wnt gene subfamilies included protein multiple-sequence alignments, phylogenetic relationship analyses (Fig. 5B), BLAST results, and occasionally protein modeling. This workflow was performed on the three represented botryllids in order to confirm and compare their Wnt repertoires. Analysis indicates that the botryllids share very similar Wnt repertoires, including extensive duplication of Wnt5. Interestingly, the botryllids lack Wnt3 that other ascidians retain. However, consistent with other ascidians, botryllids lack Wnt4 and Wnt8, but do have a WntA (Fig. 5A).

Particular attention to manual annotation was required for Wnts whose subfamily was ambiguous upon initial analysis, including Wnt1, 11, and A. As discussed with the annotation, Wnt1 was identified only in the GeneMark-ES predicted gene models. Wnt11 annotated as multiple different Wnt subfamilies; it is likely that highly conserved sequences between Wnts may have led to different subfamily categorizations when using different databases. The presence of Wnt11 in *B. violaceus* confirms the variability of Wnt11 in tunicates: absent in *B. leachii, M. occulata* and *M. occidentalis*, while present in *B. schlosseri* and other surveyed tunicates (Somorjai *et al*. 2018; Martí-Solans *et al*. 2021). Wnt11 activates both the traditionally-defined canonical and non-canonical Wnt signaling pathways; studies in *Xenopus* have found that the canonical pathway required for axis formation is activated by Wnt11, an interesting implication for further research into ascidian development (Tao *et al*. 2005; Lemaire *et al*. 2008). Initial databases used lacked the classification of WntA, so the sequence first appeared in manual annotation as a Wnt4-like gene. Phylogenetic analysis represented by Fig. 5B contained well-annotated WntA protein sequences obtained from recent literature (Martí-Solans *et al*. 2021), confirming the presence of WntA and lack of Wnt4, consistent with other tunicates.

We identified only one Wnt gene with multiple copies in the *B. violaceus* genome: four distinct Wnt5 genes (Fig. 4). Wnt5 gene duplications have previously been found in the greater order to which botryllids belong (Somorjai *et al*. 2018). The *B. violaceus* Wnt 5 genes are found in tandem, mirroring the Wnt5 gene architecture of *B. leachii* (Blanchoud *et al*. 2018; Somorjai *et al*. 2018). In addition, a potential Wnt 5 pseudogene was found between Wnt5.1 and Wnt5.2, which was similarly identified in *B. leachii*; this classification as a pseudogene is due to a smaller gene and coding sequence length when compared to other Wnt 5 genes. The parallels in the Wnt5 cluster likely indicate that the duplications and pseudogene developed in the last common ancestor of these sister species. Pseudogene conservation across species has been previously observed (Mahmudi *et al*. 2015). Additionally, our phylogeny supports independent Wnt 5 duplications after separation of the *Botrylloides* and *Molgula* genera; this pattern was previously suggested for *Molgula* and *Halocynthia* (Somorjai *et al*. 2018). Some Wnt 5 duplications are shared between the more closely related *Botrylloides* and *Halocynthia* genera, but additional duplications appear to have occurred in the botryllids (Fig. 5A). The Wnt5 gene expansion is absent in other invertebrate chordates and may be related to the unique regenerative capabilities of ascidians (Somorjai et al. 2018). Wnt5 activates the non-canonical Wnt signaling pathways, along with Wnt4, Wnt6, and Wnt11 (Schubert and Holland 2013; Komiya and Habas 2008), and non-canonical Wnt signaling has been implicated in tissue regeneration (Zondag *et al*. 2016; Hu et al. 2008). In addition, the extensive Wnt5 gene duplications may partially substitute for the functional role of Wnt4 in *B. violaceus* and *B. leachii*, and for the functional role of Wnt11 in *B. leachii* in certain contexts (Somorjai et al. 2018; Croce et al. 2006).

## Conclusions

Non-model organisms often display interesting biological processes but lack the availability of high-quality reference genomes. Our study provides a quality hybrid genome assembly and annotation for a non-model organism, *Botrylloides violaceus*. This genome adds to a growing dataset of ascidian genomes, facilitating comparative genomic analyses that can provide testable hypotheses about gene and genome evolution, as well as about the genetic changes underlying processes such as regeneration.

## Availability of Supporting Data and Materials

NCBI BioProject and BioSample identifiers are PRJNA875143 and SAMN30603932, respectively. Illumina, Inc. and Oxford Nanopore Technologies sequencing data is available on NCBI SRA, SRX17396406 and SRX17396405, respectively. *B. violaceus* genome assembly is available on NCBI, JASFYC000000000. Supporting scripts and other data generated from analysis is available on GitHub at https://github.com/calpoly-bioinf/botrylloides.

## Additional Files

**Supplementary Figure S1:** Determining optimal number of clusters, *k*, by identifying local maxima of the silhouette score.

**Supplementary Figure S2:** Pairwise correlation analysis of gene counts per orthogroup.

**Supplementary Figure S3:** Orthogroup overlap between tunicate species.

**Supplementary Figure S4:** KEGG enrichment of HOG clusters.

## Declarations

### List of Abbreviations

WBR: Whole-Body Regeneration
HOG: Hierarchical Orthogroup
HOGB: Botryllid-lineage Hierachical Orthogroup

## Author’s Contributions

J.T.S.: conceptualization, experimental optimization, whole genome sequencing and assembly, comparative genomics analysis. C.A.: genome assembly and annotation, comparative genomics analysis, Wnt family analysis. C.J. and S.W.: Wnt family analysis. P.A.: computational resources and guidance. E.L.K. conceptualization, sample acquisition, supervision. J.M.D: conceptualization, supervision. All authors contributed to writing and editing the final manuscript.

## Acknowledgements

We thank Dr. David Keeling for supplying images of *B. violaceus* colonies and Dr. Dena Block for her suggestions regarding DNA extraction.

